# Modelling the *in vitro* pre-infection dynamics of *Colletotrichum* spp. isolated from yellow passion fruit in function of environmental variables

**DOI:** 10.1101/781963

**Authors:** Karinna Vieira Chiacchio Velame, Hermes Peixoto Santos Filho, Adelise de Almeida Lima, Carlos Augusto Dórea Bragança, Francisco Ferraz Laranjeira

**Author notes:** These authors contributed equally to this work (Karinna Vieira Chiacchio Velame, Francisco Ferraz Laranjeira).

## Abstract

Brazil is the largest world producer of yellow passion fruit, but the mean yield (14.3t.ha^-1^) is less than half the potential of the crop. Part of this difference can be explained by plant health problems, including anthracnose caused by *Colletotrichum* spp. In regions with favorable climatic conditions, anthracnose can be a factor of significant yield reduction, but these regions have not yet been zoned. The objective of this study was to model the pre-infection dynamics of the fungus. The influence of temperature and photoperiod was studied on mycelia growth, sporulation and conidia germination. Mathematical models were fitted to the results and the optima for the environmental variables were estimated. The maximum mycelia growth was estimated to occur at 26.5°C. Between 24.5°C and 28.5°C the fungus grew from 95% to 100% of the estimated maximum. Temperatures below 13°C or above 34°C were harmful to mycelia growth. Temperatures over 26°C were the most favorable to sporulation while below 13°C sporulation was only 5% of the maximum. Optimum germination occurred between 25°C and 29°C with the ideal wetness period between 11h and 13h. These results can be used as a basis for zoning the risk of anthracnose occurrence in passion fruit producing regions.

**Significance and Impact of the Study:** Many diseases affect the yellow passion fruit crop, limiting its yield; among them anthracnose, caused by *Colletotrichum* spp. The disease occurs in both field (leaf and stem symptoms) and post-harvest (fruits) conditions. Understanding the role environmental conditions play in the biological cycle of such diseases is essential for developing management strategies. By modelling mycelial growth, spore production and spore germination of *Colletotrichum* spp. as affected by temperature, photoperiod and wetness period, it was possible to characterize the pathogen’s pre-infectional dynamics. The results should be used as a first approximation to estimate the risk of anthracnose occurrence in pre- or post-harvest.

## Introduction

The yellow passion fruit (*Passiflora edulis* Simns f. *flavicarpa* Deg.) is a fruit tree of key social and economic importance in Brazil. This country is the largest world producer, but Brazilian productivity is low. Pests can explain part of this productive gap, including passion fruit anthracnose, a disease caused by different species of the genus *Colletotrichum* (Damm et al., 2012). This disease affects plant development, productivity and can reduce the plantations’ life (Liberato and Costa, 2001). The fungus infects all the organs, penetrating even through the intact surface of the fruit (Benato, 1999). It is also a serious post-harvest problem (Picanço and Bruckner, 2001; Serra and Silva, 2004).

The first symptoms occur on half to fully opened leaves, on which small pale spots are observed with edges darker than the foliar surface. During colonization, white or grey areas appear in the center of the necrosis material (Yamashiro, 1987; Shoroeder et al., 1997). The lesions are elongated on the branches (Figure S1), and develop into cankers that expose the vascular tissue. The cankers become greyish, with black spots, corresponding to the fungus structures.

The lesions are depressed on the fruits, in which black fructifications of the fungus develop. Dry rot reaches the inside of the fruit causing it to wilt and fall (Santos Filho et al., 2004). The fungus survives saprophytically on crop remains or in infected tissues on the passion fruit tree itself. Raindrops and spray irrigation are the main dissemination agents (Francisco Neto, 1995).

Anthracnose control is considered difficult because few agricultural chemicals have been registered for the yellow passion fruit crop (Liberato and Costa, 2001). Thus planting in locations less favorable to the disease, associated to more resistant varieties, may be a strategy to be explored. According to Dias et al. (2005), temperature is one of the climatic variables that most influences infection and later colonization of *Colletotrichum* spp. Warm and rainy periods are generally considered conducive to greater damage, with intense leaf fall, fruit rot and die-back. According to Yamashiro (1987), temperatures between 26 °C and 28 °C, allied to intense rains favor defoliation and the disease progress.

Although there is general information on the behavior of passion fruit anthracnose, it is not sufficiently detailed to serve as risk criteria for zoning studies. Thus, our objectives were twofold (i) to model the *in vitro* dynamics of pre-infection processes of *Colletotrichum* spp. isolated from *P. edulis* f. *flavicarpa* in function of environmental variables, and (ii) develop risk criteria that could be used to establishing critical zones of disease occurrence.

## Supporting information

**Figure S1.**
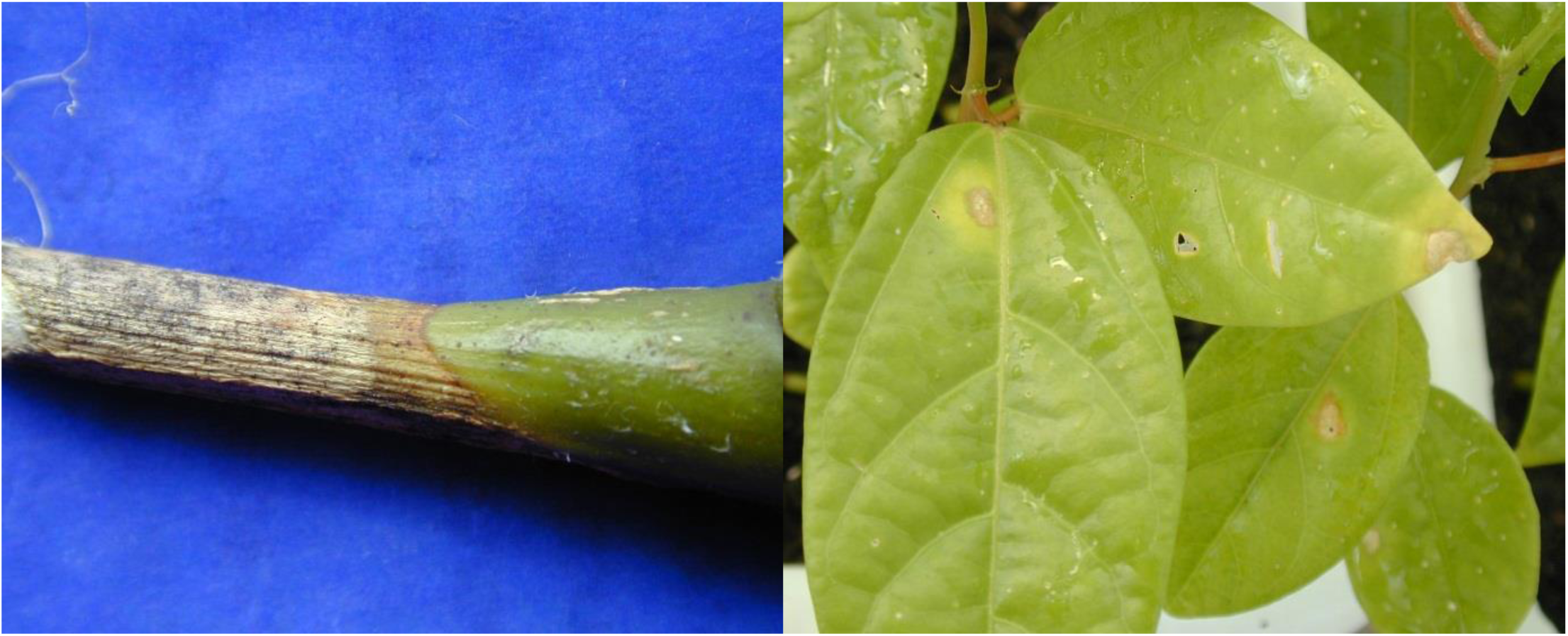
Typical twig dieback caused by *Colletotrichum* spp. in yellow passion fruit plants (left). Leaf symptoms in inoculated plants for the pathogenicity tests (right).

## Results and discussion

### Growth and sporulation in function of light period

Although the fungus grew under all the light conditions, the final mycelia area decreased when photoperiod was increased to 18 h (Figure 1A). There was a discreet increase in the mycelia area under constant light. A polynomial function (R^2^ = 0.999; without residue pattern) was fitted, with an estimated optimum of 1 h light daily (Figure 1A). Under a 12 hour light period, the mycelia growth represented 83% of that observed in constant dark. The smaller mycelia growth under the 18 hour light period was 90% of that observed under one hour of light.

**Figure 1.**
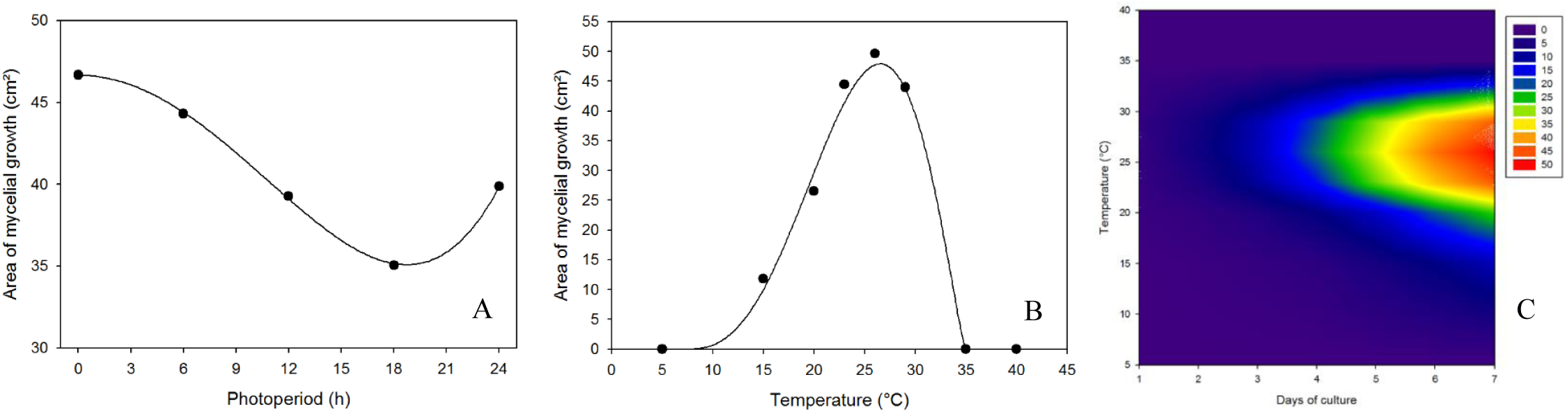
Mycelia growth of *Colletotrichum* spp. in function of environmental variables. Original data and fitted curve of grow th in function of luminosity (A). Original data and fitted curve of growth in function of temperature (B) Mycelia Growth Area = p1*((T-Tmin)^p2^)*((Tmax-T)^p3^), where p1=0.0004 Tmin=7 p2=3 Tmax=35 p3=1.3. Surface response of the mycelia growth of *Colletotrichum* spp. in function of temperature and culturing time (C). The original data are the means of two experiments and five isolates.

Light is an important environmental factor for many microorganisms and it can, for example, affect important metabolic pathways in fungus physiology (Idnurm et al., 2010). Different light intensities were reported to reduce the mycelia growth of *Colletotrichum acutatum* in the red and green light lengths (Yu et al., 2013). The negative effect of increased illumination on growth has also been reported for *Saccharomyces cerevisiae* (Roenneberg and Merrow, 2001).

This information may be useful to calibrate *in vitro* experimentation and adjust culture conditions for fungus production. These results may also have implications for studies for post-harvest fruit conservation, that is, in addition to temperature and air humidity, the light of the storage environment might influence the expansion of the lesions caused by the fungus.

The quantity of spores per dish varied very little among the light periods. Numerically the greatest sporulations were observed when six hours of daily light were used, but a model could not be found to fit the data. A simple linear regression also was not significant (P>0.76). Conidiogenesis is a complex process and can be influenced by environmental and endogenous factors (Roenneberg and Merrow, 2001; Su et al., 2012). The photoperiod can influence several aspects of fungi life cycle, such as induction or inhibition of spore formation and germination, or their growth rate (Roenneberg and Merrow, 2001). Generally, the light period is related with sporulation induction. However, light spectrum can be a limiting factor and wavelengths between 350 and 500 nm have been reported as most effective (Su et al., 2012). *Colletotrichum* spp. sporulation inhibition may have occurred due to excessive light, as reported by Leach (1964) for *Alternaria chrysanthemi*. In contrast this may have been influenced by the spectral distribution of the fluorescent light that has more frequent peaks above the range close to UV.

### Growth and sporulation in function of temperature

There was no growth at 5, 35 and 40° C (Figure 1B). The fit of the generalized beta function (R^2^ = 0.991; without residue pattern) indicated 7°C and 35°C as minimum and maximum theoretical temperatures at which growth could still occur. In spite of this, growth around these temperatures represented approximately 1% of the estimated maximum growth estimated for the optimum temperature of 26.5 °C. Curve asymmetry was not observed that could indicate a greater sensitivity to temperatures above the optimum when compared to those below the optimum. An optimum temperature range for the growth of passion fruit *Colletotrichum* spp. isolates would be between 24.5 °C and 28.5 °C. The fitted generalized beta function model indicated that between these values the fungus growth would always be between 95% and 100% of the estimated maximum. On the other hand, temperatures below 13 °C or above 34 °C would imply growth less than 10% of the estimated maximum. The calculation of the risk ranges coupled with the surface response (Figure 1C) may indicate criteria for the disease zoning. Furthermore, the risk estimates in function of decrease in temperature can be used as a guide in post-harvest passion fruit storage studies.

The mycelia growth at each temperature followed a sigmoidal pattern (Figure 1C), as the almost perfect fit indicates (Table 1). This pattern was already expected because the type of modular growth presented by fungi in general, where there is hyphal branching. However, the rate differs at each temperature, which was shown by the reduction in the number of days needed to reach 50% of the estimated maximum (parameter *b*, Table 1) as the temperature was increased.

**Table 1.**
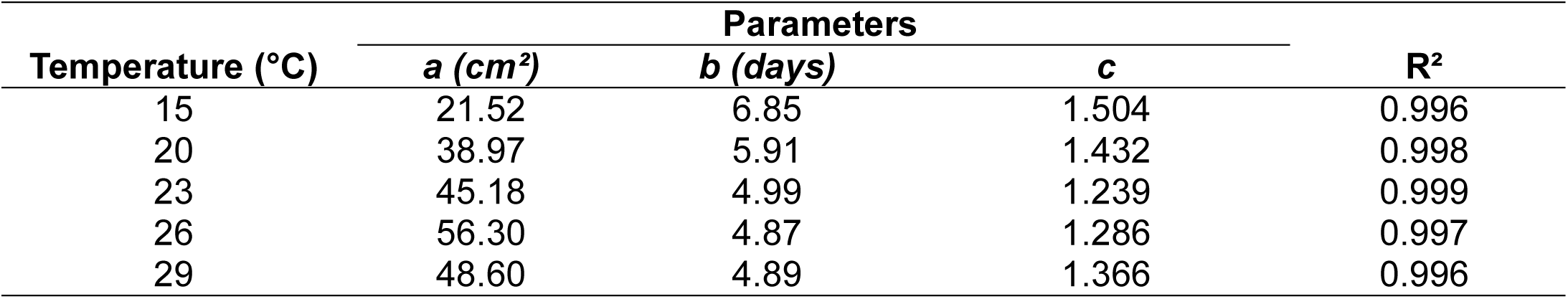
Estimated parameters and coefficient of determination of the fit of the sigmoid model ACM=a/(1+exp(-(time-b)/c))) to the progress of the area of mycelia growth of *Colletotrichum* spp. isolates from yellow passion fruit in functions of days of culture at five temperatures. The parameters *a* and *b* represent, respectively, the maximum estimated growth area and the number of days to reach 50% of the maximum growth, while *c* is a shape parameter.

Spore production in function of temperature can be explained by a beta generalized function (R^2^ = 0.979, without pattern in the residues) with an estimated optimum temperature of 29.9°C (Figure 2A). It was estimated that between 95% and 100% of the maximum spore production should occur in the range between 28.2 °C and 31.4 °C. The strong asymmetry of the fitted curve (Figure 2A) indicates that high temperatures have a more intense effect on conidia production than lower temperatures.

**Figure 2.**
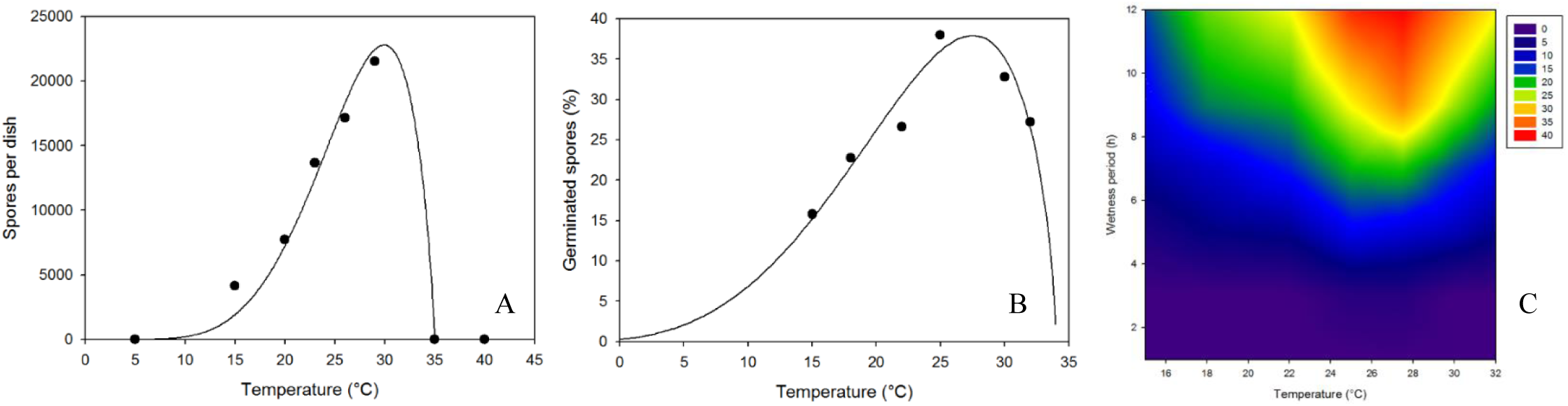
Original data and fitted curve of *Colletotrichum* spp. spore production and germination in function of temperature and wetness period. Original data and fitted curve of spore production (A) spores per dish = p1*((T - Tmin)^p2^)*((Tmax-T)^p3^)), where p1=0.00272 Tmin=3.192 p2=4.435 Tmax=35 p3= 0.842. Original data and fitted curve of the percentage of conidia germination (B) percentage of germinated spores = p1*((T-Tmin)^p2^)*((Tmax-T)^p3^)), where p1=4.2e-5 Tmin= −6.106 p2= 3.535 Tmax= 34.04 p3= 0.685. Surface response for *Colletotrichum* spp. conidia germination in function of temperature and wetness period (C). The original data are the means of two experiments and five isolates.

According to Maia et al. (2011) temperature has great influence on the development of species of the *Colletotrichum* genus both in the mycelia growth and sporulation, and conidia germination. The results presented here are to those of Cox and Irwin (1988) and Sutton (1992), who reported the optimal growth temperature for most species of *Colletotrichum* to be between 25 °C and 30 °C. Couto and Menezes (2004), for example, determined that the optimum temperature for sporulation and germination of *C. musae* conidia was between 26 °C and 28 °C.

### Conidia germination in function of temperature and the wetness period

The conidia germinated at all temperatures (Figure 2B), and it was not possible to determine the limiting minimum and maximum temperatures for germination after 12 hours of wetness. The germination percentage was not high, ranging from a little less than 40% to close to 15%. The fitted generalized beta function (R^2^ = 0.926) was used to estimate the optimum germination temperature (27.6 °C). It was estimated that between 95% and 100% of the maximum germination should occur in the range between 25 °C and 29.7 °C. The strong asymmetry of the fitted curve (Figure 2B) indicates that unlike the mycelia growth, the conidia germination was more affected by higher temperatures than by lower temperatures. Indeed, the maximum estimated temperature for germination was ∼34 °C, while temperatures as low as 5 °C would still be favorable, although limiting, to germination. These results are somehow different from those obtained by Francisco Neto et al (1994). They found that temperatures between 30 °C and 33°C were the most favourable for conidia germination of a single *Colletotrichum* isolate obtained from a different passion fruit species, *Passiflora alata*.

Using the fitted model, it was also estimated that 1% of the spores would be able to germinate at 3 °C, 5% at 8.5 °C, 10% at 12 °C and 25% at 19.5 °C. These estimates, however, still need to be tested experimentally because the values are outside the range used in our experiments.

At all the temperatures, the evolution of germination in function of the wetness period was explained by a sigmoid function (R^2^ ranging from 0.972 to 0.999 - Table 2). In this function parameter *a* corresponds to the estimate of the maximum percentage of germination, parameter *b* is a time in hours of wetness up to 50% of maximum germination and *c* is a shape parameter. At all the temperatures the maximum estimated germination was between 20% and a little less than 38%. Half of this maximum would occur between 6h30 and a little less than 8 h of wetness. The fitted sigmoid models were used to estimate optimum ranges for the wetness period. 90% of the maximum germination could be reached at any of the temperatures after wetness periods between ∼11h and ∼13h. This range corresponded to the maximum obtained experimentally. On the other hand, wetness periods as short as ∼2h would be sufficient to allow germination of 5% of the estimated maximum.

**Table 2.** Estimated parameters, hours of wetness until the occurrence of fractions of maximum estimated germination (gm) and coefficient of determination of the fit of the sigmoid model PEG=a/(1+exp(-(hours-b)/c))) to the progression of the percentage of germinated spores (peg) *Colletotrichum* spp. isolated from yellow passion fruit in function of hours of wetness period at seven temperatures. The parameters *a* and *b* represent, respectively, the maximum percentage estimated of germinated spores and the wetness hours to reach 50% maximum germination, while *c* is a shape parameter.

| Temperature (°C) | Parameters |  |  | R<sup>2</sup> | Wetness period (h) |  |
| --- | --- | --- | --- | --- | --- | --- |
|  | *a* (peg) | *b* (hours) | *c* |  | 5% gm | 95% gm |
| 15 | 27.61 | 7.81 | 1.874 | 0.988 | 2.3 | 13.3 |
| 18 | 20.18 | 6.99 | 1.611 | 0.982 | 2.2 | 11.7 |
| 22 | 24.43 | 6.82 | 1.645 | 0.981 | 2.0 | 11.7 |
| 25 | 34.33 | 6.51 | 1.783 | 0.972 | 1.3 | 11.8 |
| 27.5 | 37.43 | 6.65 | 1.587 | 0.999 | 2.0 | 11.3 |
| 30 | 32.78 | 7.63 | 1.837 | 0.987 | 2.2 | 13.0 |
| 32 | 27.60 | 7.81 | 1.874 | 0.988 | 2.3 | 13.3 |

Criteria for risk zoning were established from the surface response of the germination in function of temperature and wetting periods (Figure 2C). It was clear that the maximum risk was obtained at temperatures between 25°C and 29°C associated to wetness periods longer than 10h. Information on mean temperatures is easily available for most of the potential producing regions. On the other hand, although the potential wetness periods would be more difficult to obtain, it is clear that the association of such temperatures with rainy periods would increase the risks. These results corroborate field observations (Yamashiro, 1987), in addition to indicating a potential increase in risk for anthracnose when using spray irrigation. For *Colletotrichum* isolated from avocado, Tozze Junior (2012) determined a range between 15-30°C as favorable for conidia germination; 50% germination occurred after six hours of wetness. In the same study, after 12 hours wetness the percentage was 42% to 15°C, and greater than 50% between 20-30°C.

The results of the present study contribute to greater knowledge of the yellow passion fruit x *Colletotrichum* spp. pathosystem. Understanding the influence of environmental conditions on fungus species is an important factor for future studies. Furthermore, in the absence of zoning studies, the information from this study may be associated with geographic and climate information in a first approximation to estimate the risk of the disease occurring in determined periods and locations. This zoning could also be useful in the spatial characterization of the disease occurrence enabling better planning of management strategies. Specifically it is expected that the criteria developed here from the modelling of the pre-infection phase of *Colletotrichum* spp. isolated from yellow passion fruit can be used as an indication of the areas with the least risk for producing regions. It is certain that these criteria still need to be validated with field data or inoculation experiments. However, the validation should be part of future zoning studies, which was never the objective of the present study.

## Materials and methods

*Colletotrichum* spp. isolates were obtained from passion fruit leaves (*Passiflora edulis* Simns. f. *flavicarpa* Deg.) with typical anthracnose symptoms. Leaf fragments containing lesioned and healthy tissue were washed in water, 70% alcohol for 1 min, 1% sodium hypochlorite for 30 seconds and sterilized water. The fragments were placed in Petri dishes with synthetic PDA (potato-dextrose-agar) and incubated for seven days in a BOD at 25 °C and 12 h photoperiod.

To obtain monosporic cultures, part of the fungus colonies was placed in a test tube containing 10 ml sterilized distilled water and agitated for a few minutes to promote spore release. The initial suspension was diluted 10 times; 1 ml was collected and sown on Petri dishes with agar-agar culture medium. The dishes were placed in an incubating chamber, for nine hours at 25 °C. After this period, the surface of each dish was observed under an optical microscope. The spore of the fungus that had a germination tube developed and distant from others in the field of vision of the microscope lens was removed from the culture medium with the help of a sterile pointer and transferred to Petri dishes containing SMA culture medium (CaCO_3_-CaCl_2_-yeast-dextrose-passion fruit juice-agar). The dishes were placed in an incubating chamber at 25 °C, under a 12 h photoperiod for 15 days until the colonies grew. Later these colonies were replicated to new Petri dishes containing SMA culture medium and a work culture was obtained. A total of 113 monosporic cultures were obtained from the 35 initial isolates.

### Pathogenicity Test

An adaptation of the methodology by Dias (1990) was used. The yellow passion fruit plants were cultivated in a greenhouse in pots containing commercial substrate and obtained from seeds from the Active Germplasm Bank at Embrapa Cassava and Fruits. They were watered daily and every week 100 ml Hoagland solution was applied as nutritional supplement (Hoagland, 1950). Plants with 4 to 5 mature leaves were inoculated by spraying spore suspension (10^6^ spores.ml^-1^) on previously needle-injured leaves. Plants treated only with sterile water were used as the control. The plants were incubated in a wet chamber under transparent plastic bags for 24 hours before and 48 hours after inoculation. Symptom occurrence was assessed daily (Figure S1), and the incubation period was determined for each isolate. Five of the 113 monosporic isolates were selected from those that presented the shortest incubation period of up to 7 days (average of four plants per isolate). These five isolates were cultured in SMA culture medium for eight days in BOD at 29 °C and used in all the experiments, which were carried out twice.

### Growth and sporulation in function of temperature

Mycelia discs (5mm diameter) were transferred to Petri dishes containing SMA culture medium, each dish corresponding to a replication. A complete randomized design was used, with three replications per treatment. The dishes were incubated in BODs with temperatures kept at 5, 15, 20, 23, 26, 29, 35 and 40°C in the dark until any dish was completely covered by mycelia. The diameter of the colonies was measured daily in two perpendicular directions. The mycelia growth area was calculated considering the mean diameter and assuming the colony had a circular shape. At the end of the experiment, 2 ml water was added to each dish. The number of spores in the suspension was counted using a hemocytometer.

Models were fitted to obtained curves by non-linear regression (TableCurve 2D 5.01, Systat Software Inc.). A generalized beta function (Mycelia Growth Area = p1*((T-Tmin)^p2^)*((Tmax-T)^p3^)) was fitted to the mycelia growth area x temperature data. In this model p1, p2 and p3 are parameters, T is temperature, Tmin and Tmax are the minimum and maximum temperatures, respectively, estimated for minimal growth (Hau and Kranz, 1990). A sigmoid function (MGA = a/(1+ex p(-(time-b)/c)))) was fitted to the data of the mycelia growth area (MGA) in function of time for each temperature, where *a* is the asymptote, *b* is the number of days up to 50% of the estimated growth and *c* is a parameter used to calculate the variation of *b*. A beta function was fitted to the sporulation data in function of temperature similar to that previously described (spores per dish = p1*((T-Tmin)^p2^)*((Tmax-T)^p3^))). The R^2^ values and the presence of pattern in the residues graph were the criteria to accept or reject the models.

### Growth and sporulation in function of light period

Mycelia discs were replicated to SMA, with three replications per treatment, and each dish corresponding to a replication. The dishes were maintained in a BOD at constant 27.5 °C and 0, 6, 12, 18 or 24 h photoperiods until the diameter of any dish was covered. Growth and sporulation were measured as described in the previous experiment. Polynomial functions were fitted to both the growth and sporulation data in function of photoperiod. The acceptance criteria of the models were the same as in the temperature experiment.

### Conidia germination in function of temperature and wetness period

After eight days growth in SMA medium, spore suspensions were adjusted to 10^6^spores.ml^-1^. Four 40µl drops of the suspension were placed on three Petri dishes that were sealed with plastic film, placed in a wet chamber covered with plastic bags and incubated at 15, 18, 22, 27.5, 30 or 32 °C with wetness periods of 1, 2, 3, 5, 7, 9 or 12h. The wetness period was controlled by adding 20 µl lactoglycerol to each drop of suspension in the determined time. A completely randomized factorial design (eight temperatures, six wetness periods and four replications) was used and each replication consisted of one drop of suspension. The fraction of germinated spores was determined by counting 100 spores in each replication.

Models were fitted to the curves obtained by non-linear regression using the TableCurve 2D program. The sigmoid function previously described (germination percentage = a/(1+exp(-(wetness-b)/c)))) was fitted to the data for germination percentage x wetness hours of each temperature. Here the parameter *a* is interpreted as the maximum percentage of estimated germination for each given temperature and the parameter *b* as hours of wetness necessary for the germination of 50% of the estimated maximum. A beta function was fitted to the germination percentage in function of temperature similar to that previously described (percentage of germinated spores = p1*((T-Tmin)^p2^)*((Tmax-T)^p3^))). The R^2^ values and the presence of pattern in the residues graph were the criteria to accept or reject the models.

## Acknowledgements

This research was funded by Banco do Nordeste (Project FUP/ETENE – Fundeci N° 2001-1-103). The first and last authors are indebted to the CNPq, respectively, for their DCR and Productivity grants. We are also indebted to Mr. Francisco Paulo Souza for his technical assistance.

